# PAI-1 Interaction with Sortilin Related Receptor-1 is Required for Lung Fibrosis

**DOI:** 10.1101/2024.08.06.606812

**Authors:** Thomas H. Sisson, John J. Osterholzer, Lisa Leung, Venkatesha Basrur, Alexey Nesvizhskii, Natalya Subbotina, Mark Warnock, Daniel Torrente, Ammara Q Virk, Jeffrey C. Horowitz, Mary Migliorini, Dudley K. Strickland, Kevin K. Kim, Steven K. Huang, Daniel A. Lawrence

## Abstract

Plasminogen activator inhibitor-1 (PAI-1) has been previously shown to promote lung fibrosis via a mechanism that requires an intact vitronectin (VTN) binding site. In the present study, employing two distinct murine fibrosis models, we find that VTN is not required for PAI-1 to drive lung scarring. This result suggested the existence of a previously unrecognized profibrotic PAI-1-protein interaction involving the VTN-binding site for PAI-1. Using an unbiased proteomic approach, we identified sortilin related receptor 1 (SorlA) as the most highly enriched PAI-1 interactor in the fibrosing lung. We next investigated the role of SorlA in pulmonary fibrosis and found that SorlA deficiency protected against lung scarring in a murine model. We further show that, while VTN deficiency does not influence fibrogenesis in the presence or absence of PAI-1, SorlA is required for PAI-1 to promote scarring. These results, together with data showing increased SorlA levels in human IPF lung tissue, support a novel mechanism through which the potent profibrotic mediator PAI-1 drives lung fibrosis and implicate SorlA as a new therapeutic target in IPF treatment.

## Introduction

Diseases resulting in lung fibrosis remain challenging clinical problems. Idiopathic pulmonary fibrosis (IPF), the prototypical scarring disorder, is associated with significant morbidity and mortality^1^. Two FDA-approved drugs are now available for patients with IPF, but neither halts disease progression^1^. Therefore, more effective treatments are needed to improve the outcome of IPF and other fibrotic diseases. Plasminogen activator inhibitor-1 (PAI-1) represents an attractive therapeutic target for lung fibrosis based on data from several pre-clinical models employed in multiple laboratories^2–6^. Specifically, transgenic deletion of PAI-1 significantly attenuates fibrosis in complementary models^2,5,6^ whereas constitutive PAI-1 over-express exacerbates scarring following bleomycin injury^2^. Our studies confirm that PAI-1 promotes fibrosis not by modifying the early lung injury but rather through an activity exerted during the fibrotic phase of these models^7^. Despite the substantial evidence establishing a causal role for PAI-1, the precise mechanism by which PAI-1 promotes lung fibrosis remains unknown.

PAI-1 is a multifunctional protein with inhibitory activity against the endogenous plasminogen activators. By reducing plasmin activity, PAI-1 favors the persistence of fibrin in areas of tissue injury^5,8^. PAI-1 also binds to the somatomedin-B domain of vitronectin (VTN), a provisional matrix molecule. Adhesion to VTN enhances PAI-1 stability and disrupts cell adherence to this protein^9–13^. In prior studies, using transgenic mice and PAI-1 proteins that lack specific functions, we determined that amino acid residues necessary for VTN binding are critical for the profibrotic activity of PAI-1^7^, and we hypothesized that an interaction between PAI-1 and VTN is necessary for PAI-1-dependent fibrogenesis. We speculated that PAI-1 binding to VTN might disrupt the protective effects of this provisional matrix molecule on alveolar epithelial cell survival and wound repair, processes thought to counteract fibrosis^14,15^.

In the present study, we interrogated the importance of VTN to the profibrotic activity of PAI-1 using two distinct lung fibrosis models. Surprisingly, we found that PAI-1 promotes lung scarring in the presence or absence of VTN. This observation prompted an unbiased proteomic analysis to identify novel PAI-1 binding partners in the injured lung that might regulate its pro-fibrotic activity, and we identified Sortilin related receptor 1 (SorlA) as the most highly enriched PAI-1 interactor. SorlA is a multidomain, mosaic receptor involved in internalizing and sorting cargo proteins^16,17^. Prior studies identified roles for SorlA in human disease, but this protein has not been studied in the context of pulmonary fibrosis^18^. After identifying an interaction between PAI-1 and SorlA, we established that the binding site on PAI-1 to SorlA overlaps with its VTN binding site. Furthermore, we detected increased SorlA expression in lung tissue explants from patients with IPF, and using SorlA heterozygous and deficient mice, we established a gene-dose effect of SorlA in the development of lung fibrosis. Importantly, we confirmed that SorlA expression is necessary for PAI-1 to exert its profibrotic activity. Collectively, our data identify SorlA as a novel profibrotic cofactor for PAI-1 in the injured lung and implicate SorlA in human disease.

## Materials and Methods

### Animals

Experiments complied with institutional guidelines of the University of Michigan Committee on the Use and Care of Animals (UCUCA). Mice deficient in PAI-1 (PAI-1^Null^)^19^ and VTN (VTN^Null^)^20^ were backcrossed with C57BL/6 mice for ≥ 8 generations. Transgenic mice expressing the human diphtheria toxin receptor (DTR) off of the murine SPC promoter were generated in our laboratory on a C57BL/6 background^21^. These mice were bred with PAI-1^-/-^ mice^19^ to generate offspring that carry a single copy of the DTR gene and are PAI-1 deficient (DTR+:PAI-1^Null^). C57BL/6 mice (WT) and mice with a Sortilin related receptor 1 gene trap loss of function mutation (JAX stock #007999) were purchased from Jackson Laboratories (Bar Harbor, ME). Heterozygous SorlA breeders were crossed, and littermates were randomized to our bleomycin model.

### PAI-1 mutant proteins and recombinant SMB

Wild-type and modified PAI-1 proteins were generated in collaboration with Innovative Research (Novi MI)^22^. PAI-1_RR_ contains amino acid substitutions (Thr333Arg and Ala335Arg) within the reactive center loop that do not impair VTN-binding but abolish antiprotease activity^23^. PAI-1_AK_ contains mutations (Arg101Ala and Gln123Lys) which abolish VTN-binding but do not affect antiprotease activity^24^. The SMB domain of VTN was expressed in DS2 insect cells by Strep-tag fusion (Innovative Research; Novi, MI).

### Diphtheria Toxin (DT) Administration

Weight and age-matched diphtheria toxin receptor (DTR)-expressing mice were intraperitoneally (i.p.) injected with DT (12.5 μg/kg; Sigma Chemical, St. Louis, MO) once daily for 14 days as previously described^21^.

### Bleomycin Administration

Weight and age matched mice received oropharyngeal bleomycin (2.5 u/kg in 50 μL of sterile PBS) (Sigma Pharmaceuticals) as previously described ^7^.

### Recombinant modified PAI-1 protein administration

PAI-1^Null^:VTN^Null^ and PAI-1^Null^:SorlA^Null^ transgenic mice were intraperitoneally injected with recombinant PAI-1_RR_ or PAI-1_WT_ proteins (100 µg twice daily) for 10 days as previously described.

### Hydroxyproline assay

Hydroxyproline content of the lung was measured as previously described ^21^.

### Lung histology

Left lung was inflation-fixed at 25 cm H_2_O pressure with 10% neutral-buffered formalin, removed *en bloc*, further fixed in 10% neutral-buffered formalin overnight, and paraffin embedded. Five-micron sections were stained using hematoxylin and eosin or picrosirius red methods.

### Bronchoalveolar Lavage

BAL fluid was collected as previously described^7^.

### BAL Fluid PAI-1 Concentration Measurements

Murine total and active (plasminogen activator inhibitory activity) PAI-1 concentrations were measured in BAL fluid using a microsphere-based ELISA (Luminex) as previously described ^7^.

### Mass Spectroscopy Proteomic Analysis

Bleomycin-injured PAI-1^Null^ lungs were harvested on day 10, snap frozen, pulverized into a fine powder using a cyrogrinder (Black and Decker), and resuspended in 2.5 mL RIPA buffer (150mM NaCl, 50mM Tris, pH8.0, 1% Triton X-100, 0.5% sodium deoxycholate, 0.1% SDS) with 25 µL protease inhibitor cocktail (‘Protease Inhibitor Cocktail Set III, EDTA-Free’ from Calbiochem). Lung homogenates were centrifuged and the supernatants aliquoted. 125 µg of biotinylated PAI-1_WT_ (containing the 4-stabilizing substitutions (20)) was bound to 125 µL of streptavidin-sepharose beads (GE Healthcare). PAI-1-streptavidin-sepharose beads were added to 500 µL of lung supernatant and incubated at 37°C for 1 hour with gentle agitation. Control lung homogenates were mixed with unbound streptavidin-sepharose beads. The control and PAI-1-bound streptavidin-sepharose beads were washed, re-suspended in PBS with or without 150 µL of 12.5 mM DSP reversible cross linker (Thermoscientific), and incubated 30 minutes at room temperature. Control and PAI-1 bound streptavidin-sepharose beads were then washed with 1.2 mL of 2 M Urea (in 1x TBS) x 3, resuspended in PBS, and submitted to the mass spectroscopy core for analysis.

In the mass spectroscopy core, the beads were resuspended in 50 µl of 0.1 M ammonium bicarbonate buffer (pH∼8). Cysteines were reduced by adding 50 µl of 10 mM DTT and incubating at 45C for 30 min. Samples were cooled to room temperature and alkylation of cysteines was achieved by incubating with 65 mM 2-Chloroacetamide, under darkness, for 30 min at room temperature. An overnight digestion with 1 ug sequencing grade, modified trypsin was carried out at 37C with constant shaking in a Thermomixer. Digestion was stopped by acidification and peptides were desalted using SepPak C18 cartridges using manufacturer’s protocol (Waters). Samples were completely dried using vacufuge. Resulting peptides were dissolved in 9 µl of 0.1% formic acid and 2% acetonitrile solution and 2 µl of the peptide solution were resolved on a nano-capillary reverse phase column (Acclaim PepMap C18, 2 micron, 50 cm, ThermoScientific) using a 0.1% formic acid and 2% acetonitrile (Buffer A) and 0.1% formic acid and 95% acetonitrile (Buffer B) gradient at 300 nl/min over a period of 180 min (2-25% buffer B in 110 min, 25-40% in 20 min, 40-90% in 5 min followed by holding at 90% Buffer B for 10 min and requilibration with Buffer A for 30 min). Eluent was directly introduced into Q exactive HF mass spectrometer (Thermo Scientific, San Jose CA) using an EasySpray source. MS1 scans were acquired at 60K resolution (AGC target=3x106; max IT=50 ms). Data-dependent collision induced dissociation MS/MS spectra were acquired using Top speed method (3 seconds) following each MS1 scan (NCE ∼28%; 15K resolution; AGC target 1x105; max IT 45 ms).

Proteins were identified by searching the MS/MS data against UniProt mouse protein database (17180 entries) using Proteome Discoverer (v2.1, Thermo Scientific). Search parameters included MS1 mass tolerance of 10 ppm and fragment tolerance of 0.2 Da; two missed cleavages were allowed; carbamidimethylation of cysteine was considered fixed modification and oxidation of methionine, deamidation of asparagine and glutamine were considered as potential modifications. False discovery rate (FDR) was determined using Percolator and proteins/peptides with a FDR of ≤1% were retained for further analysis.

Spectral counts for each identified protein were represented as the log 2-fold change (FC) between PAI-1 and control samples with (+) and without (-) crosslinker. The combined fold change score for each protein was calculated as a geometric mean of the two individual fold change scores. Results are reported as Fold Change (FC) = (spectral counts in PAI-1 samples/spectral counts in control samples) which was calculated for each cross-linking condition and overall. For control samples where no spectral counts were detected, an arbitrary value of 0.2 was imputed to prevent dividing by zero.

The mass spectrometry proteomics data have been deposited to the ProteomeXchange Consortium via the PRIDE [1] partner repository with the dataset identifier PXD054196.

### Human lung tissue

Lung tissue from patients with IPF were obtained from explanted organs obtained at the time of transplant. The diagnosis of IPF was established by multidisciplinary clinical consensus prior to transplant, and all explanted lung tissue was later confirmed by pathology to demonstrate a histopathologic diagnosis of usual interstitial pneumonia, the pathologic correlate of IPF. All patients provided informed consent and the study was approved by the University of Michigan Institutional Review Board (HUM00105694). Non-fibrotic lungs were obtained from healthy controls donated by Gift of Life, Michigan that were rejected for transplantation.

### SorlA expression in human lungs

150 mg wet weight human lung tissue from IPF patients and healthy controls was homogenized in 1 mL ice cold RIPA buffer (defined previously) with complete mini protease inhibitors (Roche 11836153001). Supernatants were collected following centrifugation (10,000 x g for 20 min x 2). Protein concentrations of human lung tissue lysates were measured using a Pierce BCA Protein Assay Kit (ThermoFischer Scientific 23227). 30 µg of protein was separated on 4-15% tris-glycine gradient gels (BIO-RAD 4561084) and transferred overnight at 4°C to nitrocellulose membranes (Cytiva 10600008). Membranes were blocked at room temperature with Intercept Blocking Buffer (LI-COR 927-60001), incubated overnight at 4°C with primary antibodies: mouse anti-vinculin (2 µg/mL, Santa Cruz Biotechnology sc-25336), rabbit anti-SorLA (2 µg/mL, Abcam ab190684), or mouse anti-SMA (1:100, Invitrogen MAB-11547), and then washed with TBS 0.1% Tween-20. Membranes were stained for at room temperature with secondary antibodies: donkey anti-rabbit-800 (1:20,000, LI-COR 926-32213) or goat anti-mouse-700 (1:20000, LI-COR 926-68070), washed 3-times with TBS 0.1% Tween-20, and imaged on an Odyssey CLx (LI-COR).

### Binding of PAI-1 to SorlA in human lung tissue

Human IPF lung tissue (100 mg) was homogenized in 600 µL of ice-cold binding buffer (20 mM Tris, pH8.0, 140 mM NaCl, 10 mM CaCl2, 10% glycerol, 1% NP-40) supplemented with complete Mini, EDTA-free protease inhibitor (Roche 11836170001). Supernatants were collected following centrifugation (11,000 x g for 10 min at 4 °C x 2). 50 µL of Streptavidin Mag Sepharose beads (Cytiva 28-9857-38) were washed with binding buffer. Beads were incubated with or without 10 µg of biotinylated PAI-1_WT_ in 300 µL of binding buffer (30 min, room temperature, gentle agitation) and washed with buffer (50 mM Tris, pH7.4, 150 mM NaCl). Coated and uncoated beads were incubated at room temperature with 200 µg of sample in 500 µL of binding buffer and washed with buffer. Protein was eluted by boiling beads in 40 µL of Laemmli sample buffer (BIO-RAD 1610737) supplemented with 2-Mercaptoethanol (BIO-RAD 1610710) (5 min, 100C). Eluted protein and 50 µg of original sample (input) were analyzed by Western Blot.

### Surface plasmon resonance (SPR)

Purified full-length SorlA or its VSP10 domain (Bio-Techne) were immobilized on a CM5 sensor chip surface to the levels of 5,000 and 2500 response units respectively, using a working solution of 20 µg/ml of protein in 10 mM sodium acetate, pH 4. An additional flow cell was activated and blocked with 1.0 M ethanolamine without protein to act as a control surface. Unless otherwise stated binding experiments were performed in HBS-P buffer (0.01 M HEPES, 0.15 M NaCl, 0.005% surfactant P, 1.0 mM CaCl2, pH 7.4). Experiments were performed either on a BIAcore 3000 instrument or a Biacore 8K instrument, using a flow rate of 20 µl/min at 25 °C. Sensor chip surfaces were regenerated by 30-s injections of 10mM Glycine 20mM EDTA 0.5M NaCl pH4 at a flow rate of 30 µl/min. Equilibrium binding data for PAI-1_WT_ and PAI-1_AK_ were determined by fitting the association rates to a pseudo-first order process to obtain Req. Req was then plotted against total ligand concentration, and the data were normalized to Req/Rmax to account for different amounts of protein coupled to the surface. For the inhibition of PAI-1_WT_ binding to full length SorlA by the SMB domain of VTN, 250 nM PAI-1_WT_ was preincubated with increasing concentrations of SMB prior to SPR analysis. The data were fit to a binding isotherm using non-linear regression analysis in GraphPad Prism 10 software:

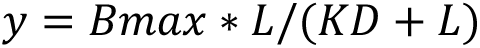

where Bmax is the Req value at saturation, L is the free ligand concentration and K*_D_* is the equilibrium binding constant.

### Statistical Analysis

Data are presented as means ± standard error of the means (SEM). For statistical analysis GraphPad Prism software was used, in any experiment with only two groups, a two-tailed t test was used. For experiments with more than two groups, a two-way ANOVA was used with a Tukey’s post hoc test for multiple comparisons. A *p* value of less than 0.05 was considered significant.

## Results

### The profibrotic effects of PAI-1 are independent of vitronectin

Previously, we found that a PAI-1 variant with intact vitronectin-binding but no protease inhibitory activity, could fully restore lung fibrosis in bleomycin-injured PAI-1^Null^ mice^7^. This result led us to hypothesize that VTN might play an antifibrotic role following lung injury, and that PAI-1 binding to VTN might abolish this protective effect. To investigate this hypothesis, we employed two complementary murine models of pulmonary fibrosis as recommended by the 2017 ATS Workshop Report^27^. One model involves selective type 2 alveolar epithelial cell (AEC2) injury via repetitive doses of diphtheria toxin (DT) administered to mice expressing the diphtheria toxin receptor (DTR) under the transcriptional control of the surfactant protein C promoter (DTR+; **Figure 1A.1**). The second model involves a single, weight-based dose of intrapulmonary bleomycin administered on day 0 with lung fibrosis assessed on day 21 (**Figure 1A.2**). In both models, as recommended by the 2017 Workshop^27^, we used hydroxyproline, a biochemical measure of collagen, to quantify the severity of lung fibrosis, and we analyzed low-power histolopathology sections to evaluate the pattern of scarring. We employed both models to interrogate the following groups of mice: 1) WT; 2) PAI-1^Null^; 3) VTN^Null^; and 4) PAI-1^Null^:VTN^Null^. In both models, untreated DTR^negative^:WT mice were used as negative controls because we found no difference in baseline lung collagen content between WT, VTN^null^, and PAI-1^null^ genotypes (**Supplemental Figure 1**). In addition, in the targeted AEC2 injury model, WT mice treated with DT were included to assess non-specific effects of DT.

**Figure 1.**
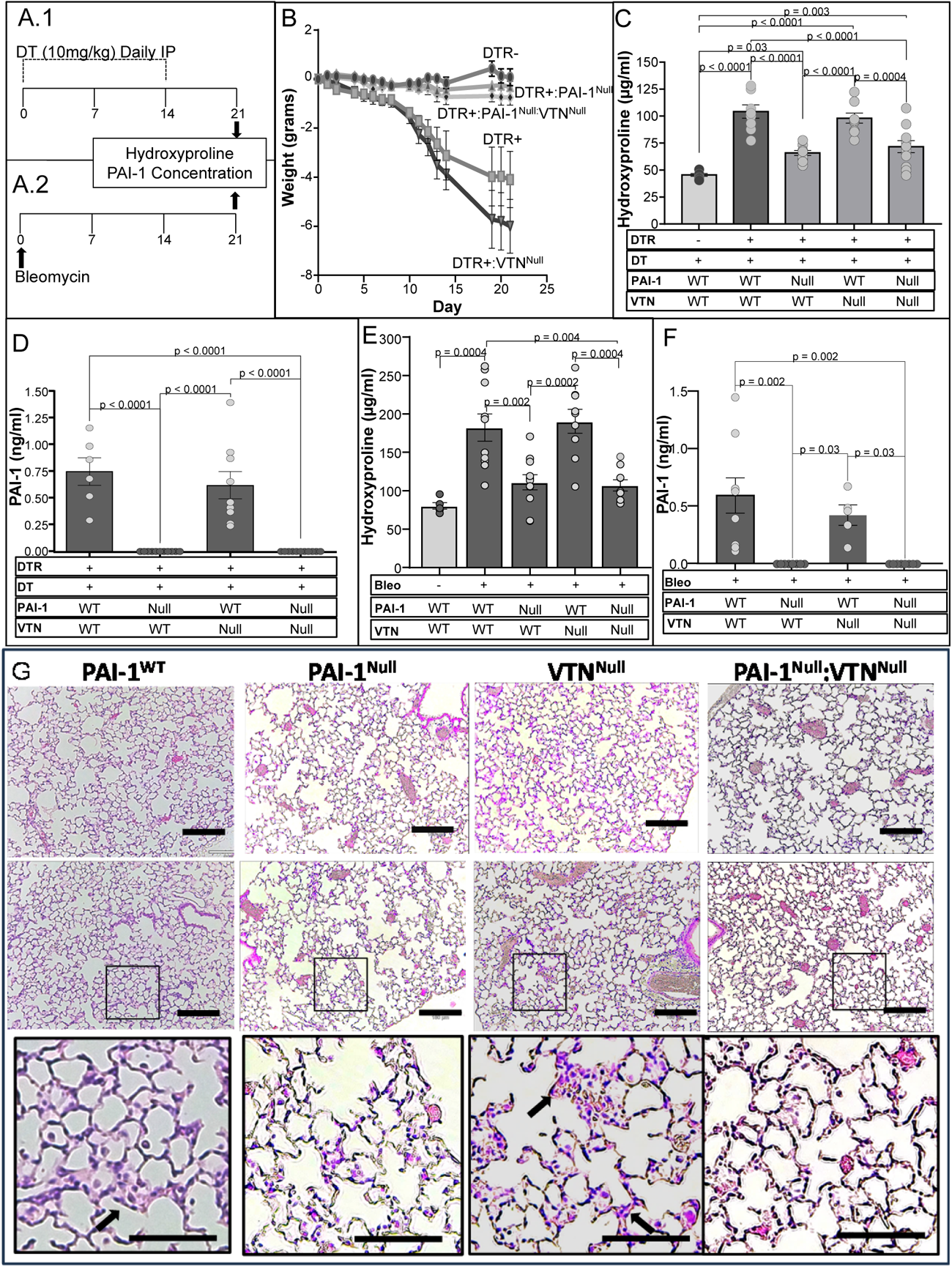
The Profibrotic Effects of PAI-1 Following Type II AEC Injury are Independent of Vitronectin. **(A.1)** Diphtheria toxin (12.5 µg/kg) was administered for 14 days to: 1) DTR^+^ mice; 2) DTR^+^:PAI-1^Null^ mice; 3) DTR^+^:VTN^Null^ mice; and 4) DTR^+^:PAI-1^Null^:VTN^Null^ mice. A control cohort of wild-type (DTR-) mice treated with DT was also included in the study protocol. (**A.2**) Bleomycin was administered (2.5 u/kg in 50 µl by the oropharyngeal route) on day 0 to: 1) wild-type mice; 2) PAI-1^Null^ mice; 3) VTN^Null^ mice; and 4) double knockout PAI-1^Null^:VTN^Null^ mice. A group of uninjured WT were included as a negative control. (**B**) In the targeted AEC2 injury model, mice were weighed at regular intervals. (**C, E**) Lungs were harvested on D21 and analyzed for hydroxyproline content. (**D, F**) Bronchoalveolar lavage samples were obtained at D21 and assayed for total PAI-1 levels. Results (B-F) are reported as the mean concentration ± SEM (n = 6-7 (B), n = 7-11 (C), n= 5-11 (D), n = 5-11 (E), and n = 5-8 (F)). Significant p values are shown from a two-way ANOVA and a Tukey’s multiple comparison test. (**G**) Different magnifications of hematoxylin and eosin-stained D21 lung sections from 2-animals in the targeted AEC2 injury model. Scale bar = 180 µm (upper panels) and 90 µm (lower panels). Arrows highlight thickened alveolar walls.

We found that DT injury to both DTR+:WT and DTR+:VTN^Null^ mice resulted in significant weight loss relative to the DT-exposed DTR^Negative^ group whose weight remained stable over the 21 day protocol (**Figure 1B**). In contrast, the DTR+:PAI-1^Null^ and the DTR+:PAI-1^Null^:VTN^Null^ mice exhibited substantially less weight loss in response to injury. Our quantification of lung fibrosis revealed that the two groups with wild-type PAI-1 expression (DTR+:WT and DTR+:VTN^Null^) developed significantly increased lung collagen content in response to DT (relative to negative control group) while the two groups deficient in PAI-1 expression (DTR+:PAI-1^Null^ and DTR+PAI-1^Null^:VTN^Null^) were protected **(Figure 1C)**. Day 21 histologic evaluation revealed evidence of interstitial thickening, increased cellularity, and enhanced picrosirius red staining in the DTR+:WT and DTR+:VTN^Null^ mice as compared to the PAI-1 deficient groups (**Figure 1G** and **Supplemental Figure 2**).

To assess for the possibility that VTN deficiency might affect PAI-1 levels in this model and thereby influence fibrogenesis, we measured PAI-1 in BAL samples obtained from each cohort of mice. Our results demonstrated no detectable PAI-1 in the PAI-1^Null^ groups and no difference in PAI-1 concentrations between the DT-injured DTR+:WT and DTR+:VTN^Null^ mice (**Figure 1D**).

In the single-dose intrapulmonary bleomycin injury model, we observed congruent results with increases in lung collagen content in injured WT and VTN^Null^ mice (relative to uninjured wild-type mice; **Figure 1E**), and attenuated fibrosis in the PAI-1-deficient groups irrespective of whether VTN was present or absent. Furthermore, the presence or absence of VTN did not affect PAI-1 levels in the BAL fluid of mice following bleomycin-induced injury (**Figure 1F**).

### Reconstitution of PAI-1 deficient mice with recombinant PAI-1 restores pulmonary fibrosis in the absence of VTN

To further rule-out a contribution of VTN in mediating the pro-fibrotic activity of PAI-1, we employed a second experimental approach. We treated injured PAI-1^Null^:VTN^Null^ mice with either PBS or PAI-1_RR_, a PAI-1 variant with intact VTN-binding activity but no plasminogen activator inhibitory activity^23^. PAI-1_RR_ was administered during the fibrotic phase from day 11 through day 21 in both models (**Figures 2A.1** and **2A.2**). This protocol was informed by previously published data in which we showed that reconstituting bleomycin-injured PAI-1^Null^ animals with PAI-1_RR_ restored lung fibrosis to a level comparable with bleomycin-injured WT animals^7^. We quantified lung fibrosis in each model with hydroxyproline and assessed the pattern of fibrosis via lung histology. For the targeted AEC2 injury model, PBS-treated DTR+:WT mice served as a negative control group while DT-treated DTR+:WT mice were included as a positive control. For the single-dose intrapulmonary bleomycin-injury model, uninjured WT mice served as a negative control while bleomycin-injured WT animals were included as a positive control. In the AEC2 injury model and consistent with data in **Figure 1C**, the DT-injured DTR+:PAI-1^Null^:VTN^Null^ group developed attenuated fibrosis when compared to the DT-injured DTR+:WT animals (**Figure 2B**). Reconstitution of the DT-injured DTR+:PAI-1^Null^:VTN^Null^ group with PAI-1_RR_ resulted in significant lung collagen accumulation that was comparable to, and not statistically different from, the DT-injured DTR+:WT group.

**Figure 2.**
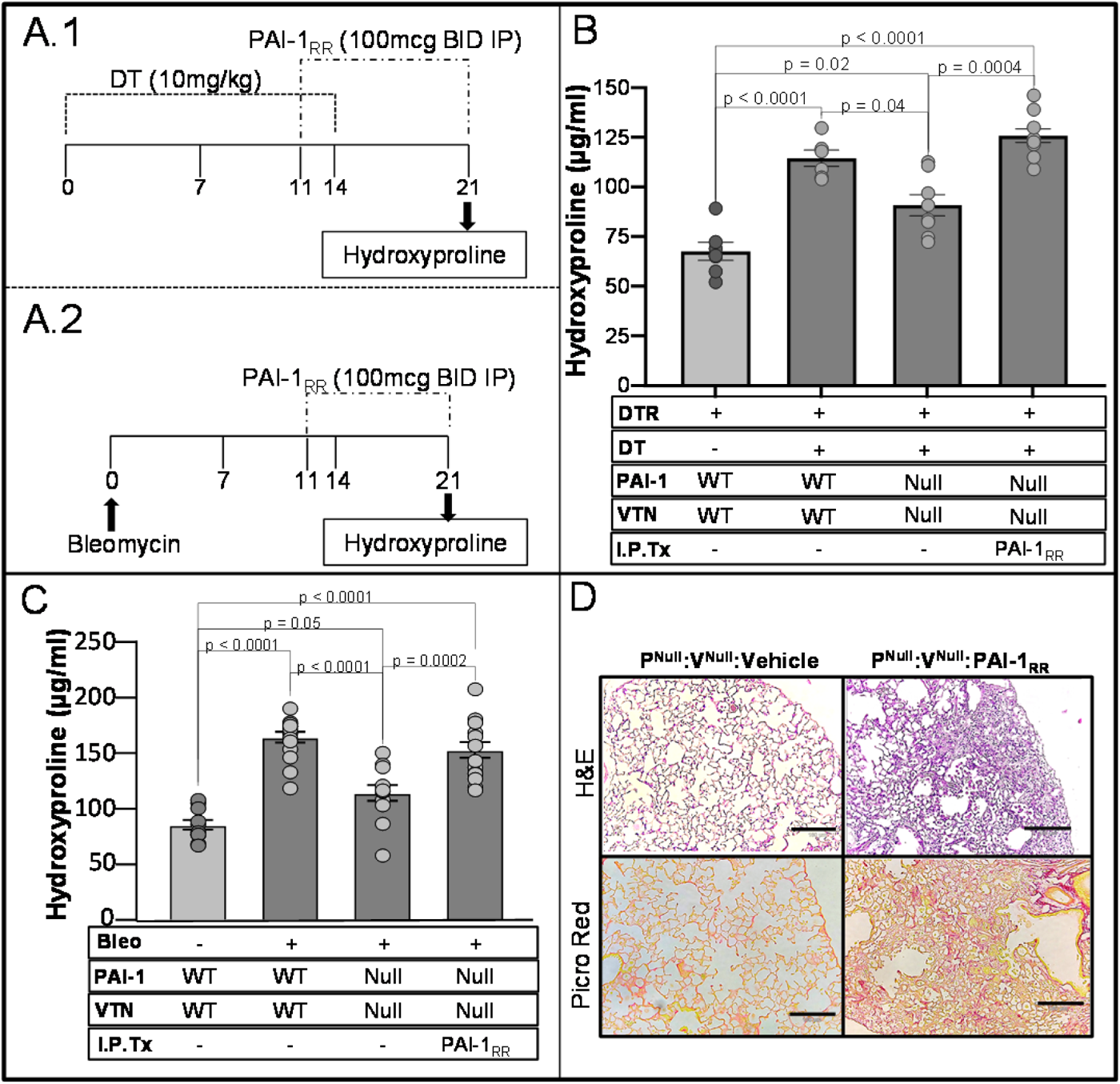
Reconstitution of PAI-1^Null^:VTN^Null^ mice with PAI-1_RR_ restores pulmonary fibrosis in a vitronectin independent manner. **(A.1)** Diphtheria toxin (10.0 µg/kg) was administered for 14 days to DTR^+^ and DTR^+^:PAI-1^Null^:VTN^Null^ mice. On D11, DT-injured DTR+:PAI-1^Null^:VTN^Null^ mice received either i.p. recombinant PAI-1_RR_ (deficient anti-protease activity but intact VTN-binding) at 100 µg twice daily or an equivalent volume of PBS. Control cohorts included DT-injured-DTR+ mice and uninjured DTR+ mice (both without PAI-1_RR_ reconstitution). **(A.2)** Bleomycin was administered (2.5 u/kg) to double knockout PAI-1^Null^:VTN^Null^ mice. Beginning on day 11, PAI-1^Null^:VTN^Null^ mice were treated with either recombinant PAI-1_RR_ or PBS. Control cohorts of mice receiving PBS in place of PAI-1_RR_ included bleomycin-injured and uninjured wild type mice. (**B, C**) Lungs were harvested on Day 21 and analyzed for hydroxyproline content. (**D**) Harvested lungs were inflating fixed, sectioned, and stained by hematoxylin and eosin (**top panels**) and picrosirius red (**bottom panels**). Scale bar = 180 µm. Results (**B, C**) are reported as the mean concentration ± SEM (n = 7-10 (B)), n = 11-15 (C)). Significant p values are shown for comparisons performed using a two-way ANOVA and a Tukey’s multiple comparison test.

In the single-dose intrapulmonary bleomycin model, we observed similar findings in that the PAI-1^Null^:VTN^Null^ group accumulated less hydroxyproline following injury when compared to the bleomycin-injured WT group. Reconstitution of PAI-1^Null^:VTN^Null^ mice with PAI-1_RR_, as in the AEC2 injury model, significantly worsen lung fibrosis as assessed by hydroxyproline (**Figure 2C**) and lung histologic analysis which revealed prominent fibrotic lesions in H&E- and picrosirius red-stained sections (**Figure 2D**).

Together, data from these complementary models demonstrate that the presence of PAI-1 during the fibrotic phase of lung injury is critical for the development of lung fibrosis <u>whether VTN is present or not</u> (**Figure 1** and **Figure 2)**. These observations support the conclusion that the pro-fibrotic function of PAI-1 is acting independently of VTN and suggest that the residues required for VTN-binding in PAI-1 likely interact with another target to exacerbate fibrosis.

### Identification of sortilin related receptor 1 as a novel PAI-1 binding partner

After finding that VTN is not required for the pro-fibrotic activity of PAI-1, we sought to identify novel PAI-1 interactors in the injured lung. To accomplish this goal, we incubated homogenized lung tissue from PAI-1^Null^ mice at day 10 post-bleomycin injury with streptavidin beads coated with biotin-tagged PAI-1. To a subset of samples, a reversible cross linker (DSP) was added to enhance the detection of weak affinity interactions. We then analyzed the abundance of proteins (spectral counts) bound to the PAI-1-coated beads relative to control beads via mass spectroscopy. Spectral counts for each identified protein were represented as the log 2-Fold Change (FC) between PAI-1 and control samples with (FC+) and without (FC-) crosslinker. Finally, we calculated an average of FC+ and FC-for each protein. As a validation of this approach, we detected within the list of the top 25 most enriched proteins both tPA and VTN (**Table 1**). In addition to these two known PAI-1 targets, the most highly enriched protein was sortilin related receptor-1 (SorlA). To substantiate binding between PAI-1 and SorlA while simultaneously determining if we could detect SorlA in human IPF lung tissue, we added PAI-1-coated streptavidin beads to homogenized explanted lung tissue obtained from three patients with end-stage fibrotic disease. The captured proteins were eluted, separated by gel electrophoresis, and immunoblotted for SorlA. With this approach, we found that PAI-1 specifically binds to SorlA in fibrotic human lung tissue (**Figure 3**).

**Figure 3:**
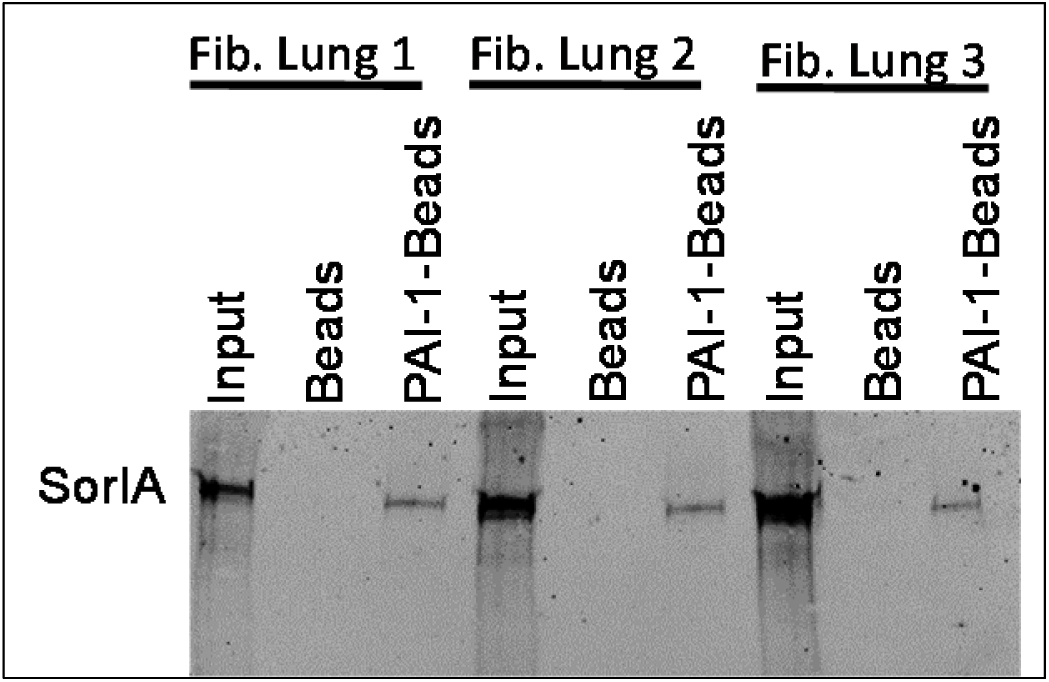
PAI-1_WT_ binds to sortlin-related receptor-1 in IPF patient lung tissue homogenates. Fibrotic lung tissue obtained from explants at the time of transplant were homogenized in binding buffer. 200 µg of each sample was incubated with either uncoated magnetic streptavidin sepharose beads or beads coated with biotin-tagged PAI-1_WT_. Beads were collected, washed, and proteins were eluted with SDS loading buffer. The initial homogenate (input) and the eluted proteins (Beads, PAI-1-Beads) were separated by SDS-PAGE, blotted, and stained with an anti-SorlA antibody. Data is displayed as a representative gel.

**Table 1.** PAI-1 binds to sortlin-related receptor-1 in the injured lung: PAI-1^Null^ mice were injured with single-dose bleomycin on D0. At D10, homogenized lung tissue was incubated with magnetic streptavidin sepharose beads coated with biotin-tagged PAI-1_WT_. A reversible cross linker (DSP) was added to a subset of these samples. Beads were collected, washed, and submitted for mass spectroscopy analysis. Spectral counts were measured for each identified protein bound to the PAI-1-coated beads, and these spectral counts were compared to the spectral counts for that same protein bound to control (uncoated) streptavidin sepharose beads. The fold-change was calculated as the log 2 of the ratio of spectral counts for PAI-1-bead/control-bead samples for each identified protein with (FC+) and without (FC-) crosslinker. The FC mean represents the geometric mean of the cross linked and non-crosslinked results. N = 3 for each condition (PAI-1-coated beads + cross-linker, PAI-1-coated beads - cross-linker, control beads + cross-linker, control beads - cross-linker).

| Protein | FC+ | FC- | FC mean |
| --- | --- | --- | --- |
| Sortilin-related receptor | 4.87 | 5.81 | 5.34 |
| Zinc finger protein 22 | 4.60 | 4.14 | 4.37 |
| Bifunctional 3'-phosphoadenosine 5'-phosphosulfate synthase 2 | 4.00 | 4.70 | 4.35 |
| ATP-dependent RNA helicase DDX42 | 4.79 | 3.54 | 4.17 |
| Polyadenylate-binding protein | 5.42 | 2.65 | 4.04 |
| BTB/POZ domain-containing adapter for CUL3-mediated RhoA degradation protein 3 | 3.46 | 4.60 | 4.03 |
| Peptidyl-prolyl cis-trans isomerase NIMA-interacting 1 | 3.66 | 4.39 | 4.03 |
| RNA binding motif protein, X-linked-like-1 | 4.92 | 2.99 | 3.96 |
| Vacuolar protein sorting-associated protein 33A | 3.66 | 4.11 | 3.89 |
| Guanylate cyclase 1, soluble, alpha 2 | 3.22 | 4.50 | 3.86 |
| Sideroflexin-2 | 3.46 | 4.27 | 3.86 |
| U4/U6.U5 tri-snRNP-associated protein 2 | 2.28 | 5.42 | 3.85 |
| Tyrosine-protein phosphatase non-receptor type 9 | 4.87 | 2.76 | 3.82 |
| Tissue-type plasminogen activator | 3.22 | 4.39 | 3.80 |
| Actin-related protein 10 | 2.58 | 4.79 | 3.68 |
| Ras GTPase-activating protein-binding protein 1 | 3.22 | 4.14 | 3.68 |
| Complement C1r-A subcomponent | 3.46 | 3.84 | 3.65 |
| Cell division cycle 5-like protein | 3.84 | 3.46 | 3.65 |
| Serine/threonine-protein kinase mTOR | 3.29 | 4.00 | 3.64 |
| Translocating chain-associated membrane protein 1 | 4.50 | 2.77 | 3.63 |
| Leucine-rich repeat transmembrane protein FLRT3 | 3.46 | 3.66 | 3.56 |
| U2 small nuclear ribonucleoprotein A' | 3.46 | 3.66 | 3.56 |
| Vitronectin | 3.52 | 3.58 | 3.55 |
| 39S ribosomal protein L23, mitochondrial | 2.94 | 4.14 | 3.54 |
| NEDD4-like E3 ubiquitin-protein ligase WWP1 | 3.22 | 3.84 | 3.53 |

To further examine PAI-1 binding to SorlA, we employed surface plasmon resonance (SPR). SorlA is a multidomain protein that contains a homologous region to the LDL receptor family of proteins, and prior studies demonstrate binding of PAI-1 to LDL receptor family members including LRP1 and SorlA^28^. SorlA also contains a vacuolar protein sorting 10 domain (VPS10) which has been shown to traffic proteins between different cytoplasmic compartments. In this experiment, we analyzed binding of PAI-1 to the isolated VPS10 domain and to full length SorlA by immobilizing both proteins on a SPR chip. Solutions containing increasing concentrations of PAI-1_WT_ and PAI-1_AK_, a mutant with intact protease inhibitory activity, no detectable affinity for VTN^24^, and attenuated profibrotic activity^7^, were flowed over the soluble forms of full length SorlA and the isolated SorlA-VPS10 domain to assess binding. As represented in **Figure 4A-B**, PAI-1_WT_ adheres 5-fold more avidly to full length SorlA than PAI-1_AK_ with respective Kd values of 126 ± 47 nM versus 540 ± 95 nM. Both PAI-1_WT_ and PAI-1_AK_ also bound to the VPS10-domain of SorlA, an interaction that has not been previously reported (**Figure 4C, D**). Like full-length SorlA, PAI-1_WT_ bound more avidly to the VPS10-domain of SorlA than PAI-1_AK_. To confirm overlap between the PAI-1 binding site on VTN and full-length SorlA, we assessed the binding of PAI-1_WT_ to SorlA in the presence of increasing concentrations of the somatomedin B (SMB) domain of VTN. We found that SMB inhibits PAI-1_WT_ binding to SorlA with an IC50 of 464 nM. This observation supports the conclusion that the binding sites on PAI-1 for VTN and SorlA overlap.

**Figure 4.**
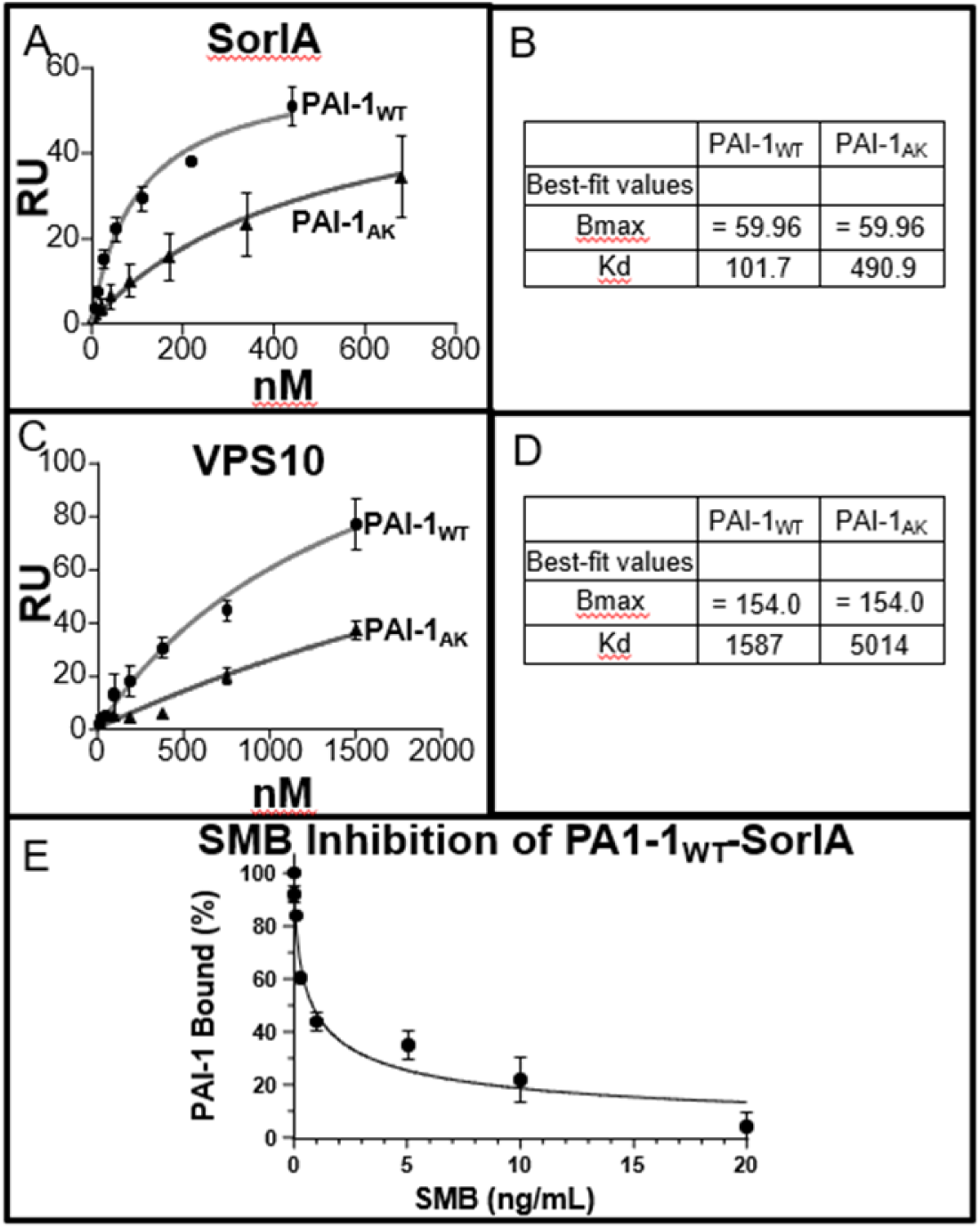
PAI-1 binds to SorlA as revealed by surface plasmon resonance analysis. Increasing concentrations of PAI-1_WT_ or PAI-1_AK_ were flowed over SPR flow cells coupled with soluble SorlA (**A, B**) or the VSP10 domain (**C, D**). Data were analyzed by equilibrium analysis by fitting the association data to a first order process to determine Req. The data were normalized to Rmax and plotted as Req/Rmax versus PAI-1 concentrations. Data were fit to a single binding site by non-linear regression analysis. Inhibition of PAI-1 (250 nM) binding to SorlA in the presence of increasing concentrations of somatomedin B (SMB) domain of VTN) **(E)**.

### Deficiency of sortilin related receptor-1 protects against lung fibrosis

After discovering that PAI-1 binds SorlA in the injured lung, we hypothesized that, if a PAI-1-SorlA interaction is required to promote pulmonary fibrosis, then SorlA deficiency should protect against scarring. To test this hypothesis, we administered single-dose bleomycin to littermate mice either deficient (SorlA^Null^), heterozygous (SorlA^Hetero^), or wild-type (SorlA^WT^) for SorlA expression. Lung collagen accumulation was quantified via hydroxyproline and the pattern of scarring was assessed by histopathology. We observed that bleomycin caused a similar initial weight loss in the three genotypes over the first 4 days of the study (**Figure 5A**). Thereafter, the SorlA^Null^ mice rapidly recovered when compared to the other two groups. Relative to the SorlA^WT^ mice, the SorlA^Hetero^ group exhibited a late recovery in weight. We found that the recovery of weight in the SorlA^Null^ and SorlA^Hetero^ groups was associated with a significant reduction in lung collagen content when compared to the SorlA^WT^ mice (**Figure 5B**). Furthermore, we observed a SorlA gene-dose response in which lung hydroxyproline content in the SorlA^Hetero^ group was intermediate between SorlA^WT^ and SorlA^Null^ mice. Histopathology revealed diffuse areas of increased lung collagen deposition and increased cellularity in SorlA^WT^ animals, and these abnormalities were much less prominent in the SorlA^Null^ mice (**Figure 5D**). We also assessed whether the SorlA expression altered PAI-1 levels in the alveolar compartment at the time of lung harvest (Day 21). We found that bleomycin injury increased PAI-1 levels in the SorlA^WT^ group compared to the control animals (**Figure 5C**). Bleomycin-injury also increased PAI-1 levels in the SorlA^Hetero^ and SorlA^Null^ mice, and there was no difference in the PAI-1 concentration relative to SorlA gene dose.

**Figure 5.**
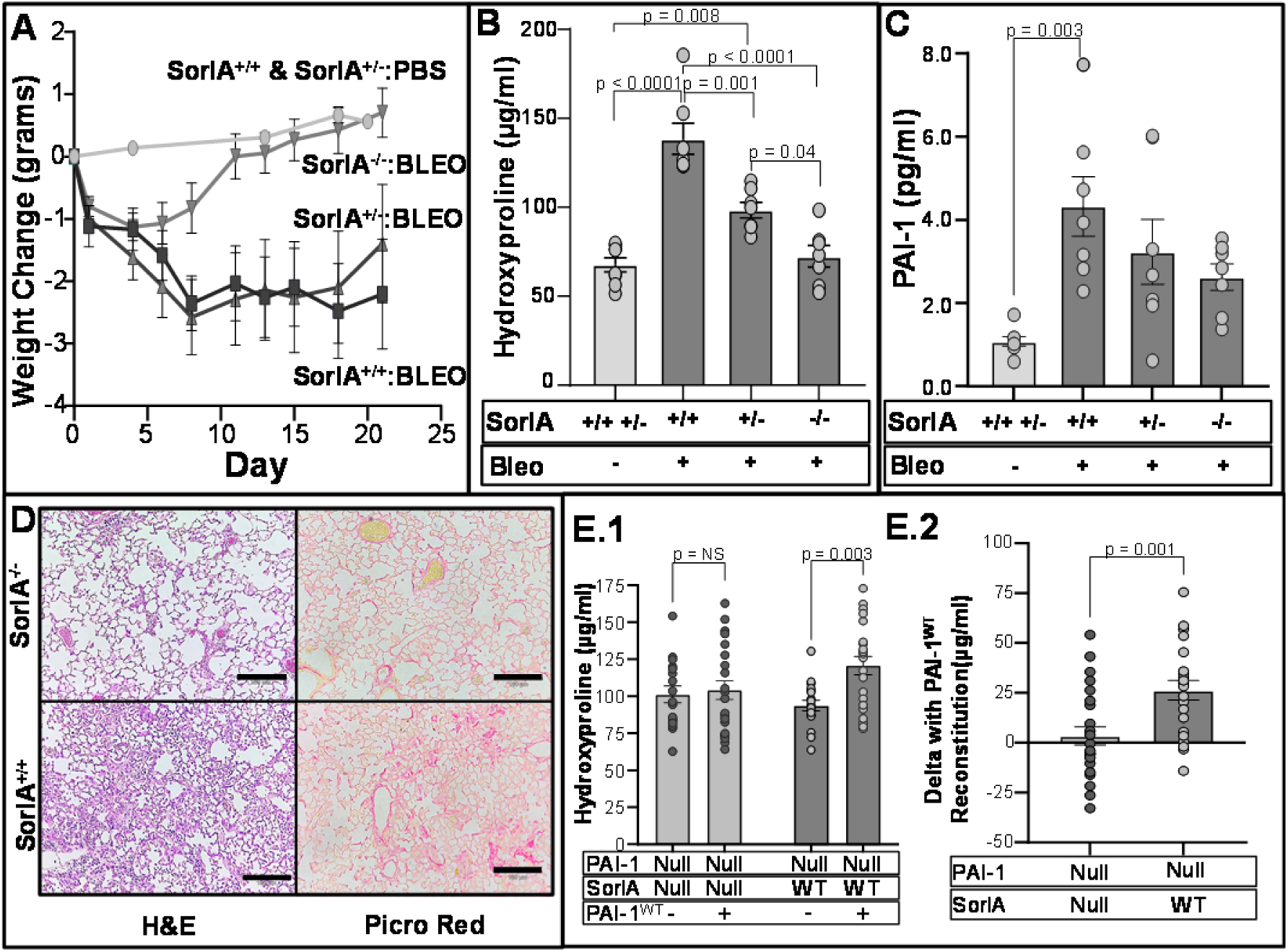
SorlA deficient and SorlA heterozygous mice are protected from fibrosis following lung injury. **(A-C)** Single dose bleomycin (2.5 u/kg in 50 µl) was administered on D0 to the following littermate cohorts: 1) SorlA^WT^, 2) SorlA^Hetero^ and 3) SorlA^Null^ mice. Control littermates (mix of SorlA^WT^, SorlA^Hetero^, SorlA^Null^) were uninjured and served to establish baseline lung collagen content. **(A)** Mice were weighed at regular intervals (n = 7-8/group) and **(B,C)** lungs and BAL were collected on D21 for hydroxyproline analysis and measurement of PAI-1 concentration (n = 7-8). Results are reported as the mean ± SEM. P values are displayed for comparisons performed using a two-way ANOVA and a Tukey’s post-hoc multiple comparison test. **(D)** Lung sections obtained at D21 were stained by H&E (left panel) and picrosirius red (right panel) and representative images are shown. Scale bar = 180 µm. **(E)** Single dose bleomycin was administered (2.5 µg/kg in 50 µl) on D0 to the following littermate cohorts: 1) PAI-1^Null^:SorlA^WT^ and 2) PAI-1^Null^:SorlA^Null^. On D11, subsets of mice from each genotype were administered either i.p. recombinant PAI-1_WT_ at 100 µg twice daily or an equivalent volume of PBS. On day 21, lungs were analyzed for hydroxyproline content. **(E.1)** Mean hydroxyproline values in each group (n = 17-24/group). **(E.2)** Delta in hydroxyproline in PAI-1Null:SorlA^Null^ treated with/without PAI-1_WT_ and PAI-1^Null^:SorlA^WT^ mice treated with/without PAI-1_WT_. Data is reported as mean ± SEM. P values are shown using a two-way ANOVA with Tukey’s multiple comparison test (B, C), grouped two-way ANOVA with Sidak Test for multiple comparisons (E.1), and an unpaired t-test (E.2). NS = non-significant.

To test whether an interaction between SorlA and PAI-1 is required for PAI-1 to exert its profibrotic activity, we administered bleomycin to PAI-1^Null^ littermate mice that were either SorlA deficient (PAI-1^Null^:SorlA^Null^) or had wild-type SorlA expression (PAI-1^Null^:SorlA^WT^). On day 11, subsets of each genotype were administered recombinant PAI-1_WT_ for 10 days and then lungs were analyzed for hydroxyproline content. We found that reconstitution of PAI-1 in the PAI-1^Null^:SorlA^WT^ group resulted in an increase in lung collagen, consistent with prior experiments (**Figure 5E.1 and E.2**, **Figure 1**, and(5)). In contrast, the administration of PAI-1_WT_ to the PAI-1^Null^:SorlA^Null^ group did not induce hydroxyproline accumulation. These results support the hypothesis that an interaction with SorlA is required for PAI-1 to express its profibrotic action.

### Sortilin related receptor 1 is increased in human fibrotic lung tissue

After identifying a critical role for SorlA in the development of fibrosis following bleomycin injury, we investigated whether SorlA is upregulated in human disease by measuring expression (via Western blotting) in lung tissue from patients with end-stage lung fibrosis (predominantly IPF) versus healthy controls. In these same samples, the expression of alpha-smooth muscle actin (αSMA; Western blotting) and active PAI-1 (ELISA) were analyzed. Compared to healthy tissue, IPF lungs exhibited increased levels of SorlA, αSMA, and active PAI-1 (**Figure 6A**). Of note, we identified a significant correlation between SorlA and αSMA in the fibrotic samples whereas no correlation between these proteins existed in healthy donors (**Supplemental Figure 4**).

**Figure 6.**
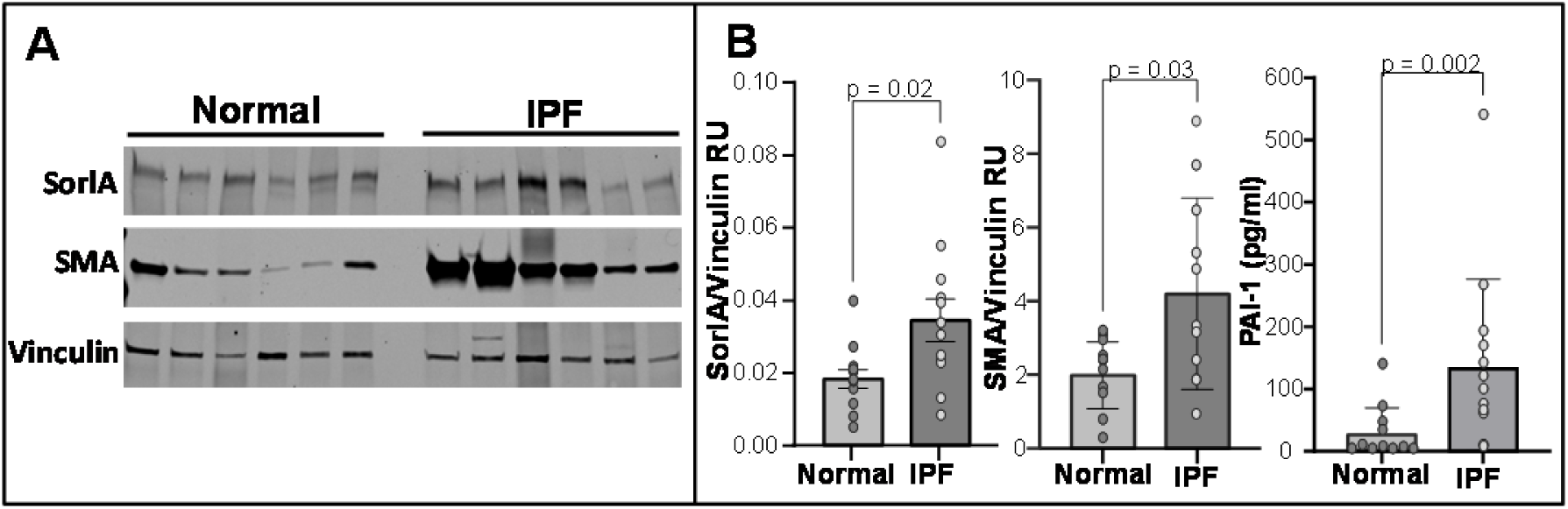
SorlA expression is increased in human IPF lung tissue: (**A**) Equal quantities of protein from homogenized IPF or normal control lung tissue was separated by SDS-PAGE (n = 6) and (**B**) analyzed by Western blot for SorlA and αSMA (normalized to vinculin, n = 13) and by ELISA for active PAI-1 (normalized to total lung protein concentration, n = 13). Data are represented as mean ± SEM. Significant p values are shown for comparisons performed using a parametric or Mann–Whitney U nonparametric two tailed t-test.

## Discussion

PAI-1 is critical to the development of tissue fibrosis in multiple organs^2,3,5,6,29–35^, and understanding its mechanism of action will shed important insight into the pathogenesis of these difficult to treat diseases. Early studies focused on the plasminogen activator inhibitory function and the preservation of fibrin in injured tissue as being essential to the profibrotic activity of PAI-1^2,36,37^. However, the demonstration that fibrin deficiency does not protect against lung fibrosis^5^, and that the tPA/uPA inhibitory activity of PAI-1 is dispensable^7^, indicated that the regulation of fibrinolysis is not the primary function whereby PAI-1 exacerbates lung scarring. Furthermore, we found that PAI-1 predominantly drives pulmonary fibrosis through a mechanism that requires its VTN-binding site^7^. Therefore, it was surprising to find in the present study that VTN is dispensable for PAI-1 to exert its profibrotic effect. This observation indicated to us that the mutations in PAI-1 that disrupt VTN-binding likely mediate interactions with other profibrotic factors. To identify novel PAI-1 interactors, we performed an unbiased proteomics analysis and discovered SorlA as the most highly enriched PAI-1 binding partner in the injured lung. This finding prompted us to investigate the role of SorlA in fibrogenesis, and we established for the first time that SorlA exacerbates lung fibrosis in a murine model and that SorlA expression is increased in human fibrotic lung tissue.

SorlA contains domains that share homology with the low-density lipoprotein receptor (LDLR) and the vacuolar protein sorting 10 (VPS10) receptor protein families. Lipoprotein receptor-related protein-1 (LRP-1) is an LDLR family member that binds PAI-1^38,39^. The homology between SorlA and LRP family members led Gliemann and colleagues to investigate an interaction between PAI-1 and SorlA^28^. They found that LRP and SorlA possess similar affinities for PAI-1, which they suggested were mediated by the ligand binding domains on the two receptors. In the present study, we extend the understanding of the PAI-1-SorA interaction by showing that PAI-1 also binds to the VPS10 domain. In addition, we found that mutations in PAI-1 that disrupt binding to VTN also reduce affinity to SorlA. We further discovered that the SMB-domain of VTN can compete PAI-1 binding to SorlA, indicating that the amino acids of PAI-1 mediating VTN-binding are also important in SorlA-binding. However, the mutations in PAI-1_AK_ that completely disrupt VTN binding do not fully abrogate binding to SorlA suggesting that other PAI-1 residues contribute to the SorlA interaction. We previously found that PAI-1_AK_ has significantly attenuated *in vivo* profibrotic activity^7^, but whether this reduced pro-fibrotic activity of PAI-1_AK_ is entirely the byproduct of impaired SorlA-binding or whether these same mutations also interfere with binding to other critical factors will require additional study. Finally, both SorlA and LRP have been shown to mediate PAI-1 intracellular uptake. The internalization of PAI-1 by LRP leads to lysosomal degradation^38^. With the known role of SorlA in protein sorting, we speculate that PAI-1 uptake by this receptor may circumvent targeting to the lysosome and allow trafficking to other cytoplasmic compartments as we recently showed for the uptake of the Tau protein^26^.

The critical role that SorlA plays in the PAI-1 profibrotic pathway is substantiated by our *in vivo* studies in SorlA-deficient and SorlA-heterozygous mice. Specifically, we identified a SorlA gene-dose effect in modulating the severity of lung scarring following bleomycin-injury. This data is remarkably congruent with that observed in PAI-1-deficient and PAI-1-heterozygous mice following the same insult^2^. We substantiated the importance of a PAI-1-SorlA interaction in lung fibrosis through a reconstitution experiment in PAI-1^Null^:SorlA^Null^ mice where administering PAI-1_WT_ following lung injury did not increase fibrosis. This result contrasts data with PAI-1 reconstitution in PAI-1^Null^:VTN^Null^ mice (**Figure 2**). Finally, the increased expression of SorlA in human IPF lung tissue lends additional credence to a role for SorlA in lung fibrogenesis.

How the SorlA-PAI-1 interaction regulates fibrogenesis will require additional studies, yet SorlA has been implicated in other aging-related disorders including cardiovascular disease and Alzheimer’s disease^40–46^. The pathogenic role of SorlA in these seemly disparate conditions leads us to speculate about shared mechanisms. SorlA is hypothesized to impact atherosclerosis pathogenesis through internalization of cell surface receptors involved in smooth muscle cell migration, and in Alzheimer’s dementia, SorlA influences the intracellular trafficking and accumulation of toxic Tau and aβ protein^26,28,47–51^. SorlA also facilitates insulin receptor localization and signaling, and SorlA deficient mice are protected from high fat feeding^52^, demonstrating a phenotype that is similar to the that of PAI-1null mice^53–56^. As an extension of these pathways, it is enticing to postulate that SorlA may control the intracellular uptake and/or sorting of PAI-1, and data from several recent reports identify intracellular functions of PAI-1^57,58^. Alternatively, SorlA could affect how PAI-1 and LRP coordinate the signaling and recycling of cell surface receptors. Ultimately, understanding the pathway(s) mediated by SorlA that exacerbate fibrosis will shed light on novel therapeutic targets. Importantly, SorlA does not appear to regulate fibrogenesis by modulating PAI-1 levels, as we found no difference in day 21-post-bleomycin BAL PAI-1 concentrations in SorlA^WT^, SorlA^Hetero^, and SorlA^Null^ mice. We acknowledge, however, that our analysis of PAI-1 at a single time point may have missed an effect of SorlA on PAI-1 at an earlier stage of scarring.

To conclude, we have further elucidated the mechanism by which PAI-1 contributes to lung fibrogenesis. We confirmed the critical role of residues in PAI-1 necessary for VTN-binding, but we discovered that VTN is dispensable in the development of fibrosis. This finding led us to discover SorlA as a highly enriched PAI-1 binding partner in the fibrosing lung, and follow-up experiments further characterized the interaction between PAI-1 and SorlA. This led us to establish a critical role for SorlA in a murine model of lung fibrosis and to determine that SorlA expression is increased in human IPF lung tissue. These novel observations provide a deeper understanding of fibrosis pathogenesis and reveal new therapeutic targets.

## Acknowledgements

We would like to acknowledge our funding sources which include the National Institutes of Health, R01-HL153056 (THS), R01-HL163870 (THS), R01-HL055374 (DAL), R01-AG074552 (DAL), R01HL141195 (JCH), and the T32 Training Grant in Lung Disease (HL007749), the Department of Veterans Affairs Merit Review Award (JJO), and the Quest For Breath Foundation (THS).

## Author Contributions

1. John J. Osterholzer assisted in planning experiments interpreting data, developing figures and was a co-primary author with Dr. Sisson.
2. Lisa Leung performed PAI-1-SorlA binding experiments in IPF lung tissue and contributed to manuscript preparation.
3. Venkatesha Basrur and Alexey Nesvizhskii assisted in designing the proteomics experiment and interpreting of the mass spectroscopy data.
4. Natalya Subbotina performed all *in vivo* experiments.
5. Ammara Virk performed the ex vivo mass spectroscopy experiments.
6. Jeffrey C. Horowitz assisted with data interpretation and manuscript preparation.
7. Mark Warnock performed western blots on human lung tissue and assays for PAI-1 concentrations and activity.
8. Daniel Torrente performed SorlA pull-down experiments with PAI-1-labeled streptavidin beads.
9. Mary Migliorini and Dudley K. Strickland designed, performed, interpreted, and helped report the plasmon surface resonance studies.
10. Kevin Kim provided baseline hydroxyproline data from uninjured VTN^null^ mice.
11. Steven Huang provided explanted human lung tissue samples from patients with advanced fibrosis and tissue from healthy controls (rejected at time of transplantation).
12. Daniel Lawrence constructed the PAI-1 mutant proteins, and he assisted in the planning of the experiments, data interpretation, and manuscript preparation.
13. Thomas H. Sisson generated the hypothesis, planned the experiments, interpreted the data, and authored the entire manuscript.

